# Resting-state alpha reactivity is reduced in Parkinson’s disease and associated with gait variability

**DOI:** 10.1101/2024.11.27.625690

**Authors:** Ellen Lirani-Silva, Layla C. S. Silva, Diego Orcioli-Silva, Victor S. Beretta, Lucas G. S. França, Daniel B. Coelho, Rodrigo Vitorio

## Abstract

**Background:** The extent to which the cholinergic system contributes to gait impairments in Parkinson’s disease (PD) remains unclear. Electroencephalography (EEG) alpha reactivity, which refers to change in alpha power over occipital electrodes upon opening the eyes, has been suggested as a marker of cholinergic function. We compared alpha reactivity between people with PD and healthy individuals and explored its potential association with gait measures.

**Methods:** Eyes-closed and eyes-open resting-state EEG data were recorded from 20 people with idiopathic PD and 19 healthy individuals with a 64-channel EEG system. Alpha reactivity was calculated as the relative change in alpha power (8-13 Hz) over occipital electrodes from eyes-closed to eyes-open. Gait spatiotemporal measures were obtained with an electronic walkway.

**Results:** Alpha reactivity was reduced in people with PD compared to healthy individuals (U = 105, p = 0.017); the rank-biserial correlation of 0.447 indicated a moderate effect size. When controlling for global cognition (Mini Mental State Examination), the group difference in alpha reactivity was no longer significant. Alpha reactivity associated with measures of gait variability only (rho = −0.437 to −0.532).

**Conclusions:** Reduced alpha reactivity in PD is driven by levels of global cognition and suggests impaired cholinergic function in PD. Reduced alpha reactivity was associated with greater gait variability, indicating a role of the cholinergic system in the mechanisms underlying gait variability. Therefore, the cholinergic system may represent a target for treatments aiming to reduce gait variability and alpha reactivity should be further explored as an endpoint for clinical trials.

## INTRODUCTION

A prominent feature of Parkinson’s disease (PD) is impaired gait, which is marked by slow walk, short steps, increased variability and reduced automaticity.^1^ Gait automaticity refers to the ability to walk with minimal conscious effort (or executive-attentional resources).^2,3^ In PD, this automatic control is disrupted by the degenerative process that affects the basal ganglia and networks involved in automaticity. It is suggested that people with PD may compensate for deficits in automaticity by engaging cognitive resources for the control of gait,^4–7^ a process thought to be mediated by the cholinergic system.^8^ However, the levels of acetylcholine are reduced in people with PD,^9,10^ limiting the ability of the compensatory mechanism in maintaining healthy gait.

Measuring the integrity of the cholinergic system may provide valuable insights into the extent of cholinergic involvement in PD-related gait impairments. Three main techniques can assess the cholinergic system. First, molecular imaging techniques, such as single-photon emission computerized tomography and positron emission tomography, allow the assessment of cholinergic markers (presynaptic, postsynaptic and muscarinic acetylcholine receptors). Molecular imaging studies show that gait impairments relate to cortical levels of acetylcholine whereas fall history is linked to thalamic levels of acetylcholine in people with PD.^9,11–15^ In turn, freezing of gait has been linked to reduced vesicular acetylcholine transporter binding in striatal cholinergic interneurons and the limbic archicortex.^15^ Second, short latency afferent inhibition (SAI) measured with transcranial magnetic stimulation (paired with electrical stimulation of a peripheral nerve) can quantify cortical cholinergic activity.^16^ SAI studies show that lower levels of acetylcholine are associated with shortened strides, reduced gait speed and increased dual-task cost and falls in people with PD.^17,18^ Although molecular imaging and SAI studies have contributed to the understanding of the cholinergic system, such techniques involve expensive equipment, injection of tracers and/or magnetic and electric stimulation that may be uncomfortable. Such inherent characteristics may limit the utilization of molecular imaging and SAI.

The third technique for the assessment of the cholinergic system refers to electroencephalography (EEG) alpha reactivity. The concept arose from studies linking EEG alpha activity with cognitive processes, particularly attention and alertness, which are modulated by cholinergic activity. During resting-state eyes-closed EEG, a prominent rhythm in the alpha frequency band (8-13 Hz) can be observed.^19^ In healthy individuals, opening of the eyes results in marked attenuation of alpha power.^19,20^ Thus, alpha reactivity is defined as the magnitude of such attenuation (or change) in alpha power from eyes-closed to eyes-open condition, particularly in occipital electrodes. Wan *et al.*^21^ demonstrated that alpha reactivity is associated with the functional connectivity between the nucleus basalis of Meynert (a major source of cortical cholinergic innervation) and the visual cortex. Further, reduced alpha reactivity is associated with smaller volumes of the nucleus basalis of Meynert.^22^ Therefore, alpha reactivity has been suggested as a marker of cholinergic system integrity,^21,23^ with reduced alpha reactivity reflecting impaired cholinergic function.

Measuring alpha reactivity offers a novel, non-invasive and relatively inexpensive approach to explore the relationship between cholinergic system integrity and gait impairments in PD. To date, two studies have reported reduced alpha reactivity in PD^24,25^ and no previous study has investigated the association between alpha reactivity and gait in PD. Therefore, the aim of this study was to compare alpha reactivity between people with PD and healthy individuals and explore its association with gait measures representing various gait domains. By examining the link between a cholinergic marker (i.e., alpha reactivity) and gait performance, this study may provide insights into cholinergic mechanisms underlying gait impairments in PD. We hypothesise that people with PD will demonstrate reduced alpha reactivity, which will be associated with gait impairments.

## METHODS

### Participants, ethics and clinical assessment

This is an observational, cross-sectional study. Thirty-nine participants were recruited from our lab database (Sao Paulo State University, Rio Claro, Brazil), including 20 people with PD and 19 healthy individuals. All participants gave their written informed consent before they participated in the study. The study procedures were conducted in accordance with the Declaration of Helsinki, and the study protocol was approved by the local Ethics Committee (#39844814.5.0000.5465).

Patients were selected on the criteria of having a confirmed diagnosis of idiopathic PD (according to UK Parkinson’s Disease Society Brain Bank criteria). The following inclusion criteria were common for both groups: able to walk unaided and be community-dwelling. Exclusion criteria were the following: clinical diagnosis of dementia or other severe cognitive impairment (according to recommendations for utilization of the Mini-Mental State Examination in Brazil; cut-off = 20/24 points for illiterates and those who attended formal education, respectively^26^); diagnosed major depressive disorder; chronic musculoskeletal or neurological disease (other than PD). An initial anamnesis was carried out to obtain demographic information (e.g., age, height, body mass, etc.). The Unified Parkinson’s Disease Rating Scale (UPDRS) and the Hoehn and Yahr Rating Scale (H&Y) were applied to assess disease severity and disease stage, respectively. People with PD were tested in the ON state of medication (approximately one hour after having taken a dose).

### EEG recordings and processing

Resting state EEG recordings were acquired using a mobile 64-channel system (eego^TM^ sports, ANT Neuro, Enschede, The Netherlands), sampled at 1024 Hz. Waveguard caps, comprising sintered Ag/AgCl electrodes, were placed according to the 10–10 system. One channel was used to record electrooculography data, with the electrode positioned below the left eye. Participants remained seated on a comfortable chair (in a quiet room) and EEG signals were recorded for eyes-closed (5 minutes) and eyes-open (2 minutes) conditions (Figure 1). Participants were instructed to remain awake. During the eyes-open condition, participants were asked to fixate their gaze at a target placed on the wall in front of them (2m away from the chair), at the height of their eyes. The research team supervised participants during the recording to monitor adherence to the protocol.

Pre-processing steps of EEG data were performed in MATLAB (Mathworks, Natick, MA, USA), using scripts and functions based on EEGLAB 2024 (http://www.sccn.ucsd.edu/eeglab).^27^ The EEG data were pre-processed using the following pipeline. First, signals were band-pass filtered between 0.5 and 100 Hz using a second-order Butterworth filter. Line noise was attenuated with the CleanLine function (pop_cleanline) with a 60 Hz notch filter, bandwidth of 2 Hz, and a time window of 4 seconds. Noisy channels and artifactual segments were removed using the Clean Rawdata function (pop_clean_rawdata), applying a flatline threshold of 5 seconds, a channel correlation criterion of 0.8, and a burst criterion of 20 to identify and reject non-brain activity. Channels were re-referenced to the average reference. Independent component analysis (ICA) was performed using the extended version of the RUNICA algorithm (pop_runica) to decompose the data into independent components. The resulting components were classified using ICLabel (pop_iclabel), and components representing non-neural sources (e.g., eye movements, muscle artifacts)^28^ were identified with a probability threshold of 0.6 and flagged using pop_icflag. These flagged components were then removed (pop_subcomp). Finally, any previously removed channels were interpolated using spherical spline interpolation to restore the channel configuration (pop_interp). This comprehensive pipeline ensures the data are clean, free of artifacts, and ready for further analysis.

Data from three occipital electrodes (O1, O2, and Oz) were selected for the alpha reactivity analysis.^21,22^ The power spectral density (PSD) was estimated for each electrode separately, using Welch’s method with a frequency resolution of 0.25 Hz and a Hamming window. The resulting PSD for the three electrodes was averaged across the three electrodes for each condition separately (eyes-closed and eyes-open). Alpha reactivity (*α_R_*) was then calculated according to the following formula:

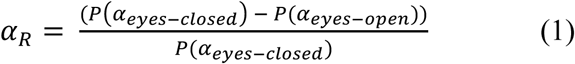

Alpha power (*P*) was computed as the relative power within 8-13Hz frequency band (normalized by the total power across all frequency bands).

### Gait assessment

Participants walked, at their preferred pace, for one minute around a 25.8 m oval circuit (with two 6.5 m parallel straights; Figure 1). Four trials/repetitions were performed. Spatiotemporal gait parameters were recorded by a 5.74 m electronic walkway (200 Hz; GAITRite®, CIR Systems, Inc., Franklin, NJ, USA) placed over the central area of the first straight segment of the circuit. We recorded an average of 107.3 [range: 97 – 127] steps for people with PD and 103.4 [range: 86 – 119] steps for healthy individuals, which is far above the minimum of 30 steps recommended for reliable measures of gait variability.^29^ A customized MATLAB algorithm was used to obtain the following gait parameters (from the GAITRite output): step length, step duration, step velocity, step width, stance time, swing time (mean of the recorded steps) and step-to-step variability in the same parameters (standard deviation of the recorded steps).

### Statistical analysis

The statistical analysis was carried out in SPSS version 26 (The International Business Machines Corporation, USA); the significance level was set to 0.05. For demographic data, unrelated sample Student’s t-tests, Mann–Whitney, and chi-square tests were employed for between-group comparisons. The Shapiro-Wilk’s test showed that alpha reactivity had non-normal distribution in our sample. Therefore, the Mann-Whitney U test was used to compare alpha reactivity between groups. The rank-biserial correlation was calculated as a measure of effect size and interpreted as: small effect (<0.3), moderate effect (0.3 to 0.5) or large effect (>0.5). Because the Mini Mental differed between groups, we used rank analysis of covariance (Quade’s test) to compare alpha reactivity between people with PD and healthy individuals while controlling for Mini Mental scores. Furthermore, the associations between alpha reactivity and gait measures or clinical scales (i.e., UPDRS, H&Y and Mini Mental) were explored using Spearman correlation coefficients.

### Data availability

Derived data supporting the findings of this study are available from the corresponding author on request.

## RESULTS

### Participants characteristics

The two groups were not significantly different in sex, age, body mass or height (Table 1). People with PD presented H&Y stages I to III, with mild to moderate disease severity (Table 1) as measured by the UPDRS. Despite having preserved global cognitive function, people with PD had lower Mini Mental scores than healthy individuals (Table 1). Relative to healthy individuals, people with PD walked with shorter and slower steps and increased step-to-step variability (Supplementary Table 1).

**Table 1.** Demographic characteristics of both groups.

| Variables | Parkinson | Control | p-value |
| --- | --- | --- | --- |
| Sex (male/female) | 8/12 | 7/12 | 0.839 |
| Age (years) | 70.6 (9.6) | 70.1 (5.6) | 0.860 |
| Body mass (kg) | 69.5 (11.5) | 68.7 (10.5) | 0.812 |
| Height (cm) | 161.2 (7.8) | 160.7 (7.3) | 0.858 |
| Mini-Mental (0-30 score) | 27.2 (1.7) | 28.8 (1.0) | 0.005* |
| UPDRS I (0-16 score) | 3.3 (1.2) | -- | -- |
| UPDRS II (0-52 score) | 9.1 (4.4) | -- | -- |
| UPDRS III (0-108 score) | 23.9 (8.9) | -- | -- |
| Hoehn & Yahr [1/1.5/2/2.5/3] | 2/4/9/4/1 | -- | -- |

\*significant difference between groups; UPDRS, Unified Parkinson's Disease Rating Scale.

### Alpha reactivity: Healthy vs PD

Figure 2 shows the power spectral density curves of the three occipital electrodes (O1, O2, and Oz) in the eyes-closed (black line) and eyes-open (red line) conditions. Alpha reactivity from eyes-closed to eyes-open condition was reduced in people with PD compared to healthy individuals (U = 105, p = 0.017, Figure 3); the rank-biserial correlation of 0.447 indicated a moderate effect size. However, when controlling for scores in Mini Mental, the group difference was no longer significant (p = 0.216, Figure 3).

### Associations with gait, cognition and clinical scales

Across the entire sample, alpha reactivity associated with measures of gait variability only (Table 2), namely: stance time variability (rho = −0.532), step velocity variability (rho = −0.515), step time variability (rho = −0.463), step length variability (rho = −0.449) and swing time variability (rho = −0.437). When only people with PD were considered for correlations, alpha reactivity also associated with measures of gait variability only (Table 2), namely: stance time variability (rho = −0.638) and swing time variability (rho = −0.544). Figure 4 illustrates the significant association between alpha reactivity and gait variability. Alpha reactivity also associated with Mini Mental (rho_sample_ = 0.346, rho_PD_ = 0.536; Table 2). Alpha reactivity was not significantly associated with other gait measures, UPDRS or H&Y scale (p > 0.05; Table 2).

**Table 2.** Spearman correlation coefficient and p-values for correlations between alpha reactivity and gait measures or clinical scales across the overall sample and separately for people with PD.

| Outcome measures | Overall (n=39) | Parkinson (n=20) |
| --- | --- | --- |
| Step velocity | $\rho = 0.289, p = 0.075$ | $\rho = 0.117, p = 0.622$ |
| Step length | $\rho = 0.294, p = 0.069$ | $\rho = 0.161, p = 0.498$ |
| Step time | $\rho = -0.078, p = 0.635$ | $\rho = 0.014, p = 0.952$ |
| Swing time | $\rho = 0.125, p = 0.450$ | $\rho = 0.271, p = 0.248$ |
| Stance time | $\rho = -0.148, p = 0.369$ | $\rho = -0.056, p = 0.816$ |
| Step velocity variability | $\rho = -0.515, p < 0.001^{**}$ | $\rho = -0.435, p = 0.056$ |
| Step length variability | $\rho = -0.449, p = 0.004^{**}$ | $\rho = -0.260, p = 0.268$ |
| Step time variability | $\rho = -0.463, p = 0.003^{**}$ | $\rho = -0.357, p = 0.122$ |
| Swing time variability | $\rho = -0.437, p = 0.005^{**}$ | $\rho = -0.544, p = 0.013^{*}$ |
| Stance time variability | $\rho = -0.532, p < 0.001^{**}$ | $\rho = -0.638, p = 0.002^{**}$ |
| Mini Mental | $\rho = 0.346, p = 0.033^{*}$ | $\rho = 0.536, p = 0.015^{*}$ |
| Hoehn & Yahr stage | -- | $\rho = 0.289, p = 0.075$ |
| UPDRS-I | -- | $\rho = 0.125, p = 0.599$ |
| UPDRS-II | -- | $\rho = -0.062, p = 0.794$ |
| UPDRS-III | -- | $\rho = -0.102, p = 0.669$ |

\*\* Significant correlation at $p < 0.01$ ;
\* Significant correlation at $p < 0.05$ .

## DISCUSSION

Alpha reactivity from eyes-closed to eyes-open condition has been suggested as a marker of cholinergic activity.^21,22,24^ We compared alpha reactivity (from occipital electrodes) between people with PD and healthy individuals. This is the first study to examine the association between alpha reactivity and gait in people with PD. We demonstrated that alpha reactivity is reduced in people with PD relative to healthy individuals. This finding confirmed our hypothesis and was largely expected, as reduced alpha reactivity in PD had been previously reported,^24,25^ and other techniques (i.e., molecular imaging and SAI) had demonstrated that cholinergic deficits are also part of pathophysiology in PD.^9,10,17,18^ However, the group difference in alpha reactivity was not significant while controlling for Mini Mental scores; and reduced alpha reactivity associated with worse Mini Mental score, indicating that the group difference in alpha reactivity is driven by levels of global cognitive function and not specific to PD. Finally, we observed that alpha reactivity was specifically associated with measures of gait variability, suggesting that cholinergic activity plays a role in the control of gait variability, with reduced cholinergic activity being associated with greater gait variability.

### Cholinergic deficits are associated with worse gait

Studies combining gait assessment and different techniques to assess cholinergic function (namely molecular imaging and SAI) have highlighted the role of the cholinergic system in gait impairments in PD. Bonnen *et al*.^11^ showed that gait speed while OFF medication was slower in people with PD with low levels of neocortical acetylcholine ([(11)C]methyl-4-piperidinyl propionate acetylcholinesterase PET imaging), but not in those with PD with normal-range levels of acetylcholine. Rochester *et al*.^18^ observed that worse SAI was associated with slower gait speed and shorter step length (while ON medication). Pelosin *et al*.^17^ observed that worse SAI is associated with increased dual-task cost of gait speed (ON medication). In contrast, we observed that reduced alpha reactivity is specifically associated with greater gait variability (but not with other gait measures). It is worth noting that our findings are in line with findings from Henderson *et al.*^30^, who observed reduced step time variability in response to rivastigmine. In combination, these findings suggest that different measures of cholinergic function (which may represent distinct sources of cholinergic activity or pathways) may selectively influence distinct aspects of gait. A larger study combining all three techniques of cholinergic function assessment with gait analysis is needed to clarify this interpretation. To date, d’Angremont *et al.*^31^ recently reported that SAI did not correlate with [^18^F]FEOBV-PET in people with PD patients (in the primary motor and somatosensory cortices and the thalamus), suggesting that SAI and [^18^F]FEOBV-PET assess different aspects of cholinergic transmission.

Alpha reactivity reflects cortical activation changes associated with attention, cognitive engagement and visual input, which are modulated by cholinergic activity.^8^ Alpha rhythms are generated in part by thalamocortical circuits, which are also modulated by acetylcholine. Cholinergic inputs to the thalamus and cortex help regulate sensory processing and attentional shifts, leading to alpha desynchronization during wakeful, eyes-open states. Further, alpha reactivity is associated with smaller volumes of the nucleus basalis of Meinert and its functional connectivity with the visual cortex.^21,22^ Thus, the specific association of alpha reactivity with gait variability may be due to gait variability being more sensitive (than other gait measures) to changes in cholinergic function within pathways involving the above-mentioned areas (i.e., thalamus, cerebral cortex and nucleus basalis of Meinert).

### Alpha reactivity is reduced in PD, but it is not specific to PD

Reduced alpha reactivity does not seem to be specific to PD. Our findings of reduced alpha reactivity in PD are in line with two recent studies.^24,25^ However, the group difference disappeared while controlling the comparison for Mini Mental scores. In addition, reduced alpha reactivity has been reported in other neurological cohorts (e.g., Alzheimer’s disease, Lewis Body Dementia, PD Dementia) compared to healthy individuals.^22,32,33^ Thus, alpha reactivity, as a stand-alone measure, does not seem to represent a marker for a specific pathology. Instead, alpha reactivity seems to be a good marker for overall integrity of the cholinergic system or cholinergic activity. This is supported by studies reporting that reduced alpha reactivity is related to a loss of cholinergic drive from the nucleus basalis of Meynert,^21,22^ a major source of cortical cholinergic innervation. Because reduced alpha reactivity has also been observed in people with Mild Cognitive Impairment, alpha reactivity should be further investigated as a potential marker for early cognitive decline.

### Clinical implications and future directions

Alpha reactivity may represent a non-invasive and relatively inexpensive option to assess cholinergic function in clinical trials involving interventions aimed to enhance cholinergic function and/or reduce gait variability. The interplay between alpha reactivity, different gait measures/domains and different cognitive domains must be further investigated in PD (as we only explored global cognition) and other neurological cohorts with gait and/or cognitive impairments.

An optimal protocol for calculating alpha reactivity must be established, with definitions of minimal number of EEG channels needed and optimal duration of the recordings and automated preprocessing steps. Thus, researchers in the field of gait research are encouraged to explore the development of alpha reactivity as a biomarker for understanding the cholinergic mechanisms associated with gait impairments and response to interventions. For example, although cholinergic augmentation with acetylcholinesterase inhibitors can improve gait and turning in PD,^34,35^ their side effects are burdensome and particularly problematic in the older adult population.^30^ Therefore, alternative treatment options to increase cholinergic function need to be investigated, including exercise, vagus nerve stimulation, transcranial direct current stimulation (tDCS) and others. To date, among various gait domains assessed in our recent study,^36^ only gait variability (i.e., step time variability) responded to the combination of aerobic exercise with anodal tDCS over the prefrontal cortex. Thus, the addition of anodal tDCS over the prefrontal cortex to a session of aerobic exercise may have increased cholinergic activity, leading to reduced gait variability.

### Limitations

A limitation of the current study is that people with PD were only tested in their ON state of dopaminergic medication, which may influence EEG recordings by increasing resting-state eyes-closed alpha power.^37^ Furthermore, our study did not assess various distinct cognitive domains, limiting the interpretations to global cognition as assessed with the Mini Mental. Given the promising nature of current findings, future studies are encouraged to address these limitations.

### Conclusion

Alpha reactivity from resting-state eyes-closed to eyes-open EEG is reduced in people with PD, suggesting impaired cholinergic function in PD. The reduced alpha reactivity is driven by the association with levels of global cognition. Furthermore, reduced alpha reactivity associated with greater gait variability, indicating a crucial role of the cholinergic system in the mechanisms underlying increased gait variability. Therefore, the cholinergic system may represent a target for treatments aiming to reduce gait variability and alpha reactivity should be further explored as a potential endpoint for clinical trials.

## ACKNOWLEDGEMENTS

Authors acknowledge participants for donating their time.

## AUTHOR CONTRIBUTIONS

## STATEMENTS AND DECLARATIONS

### Ethical considerations

The study procedures were conducted in accordance with the Declaration of Helsinki, and the study protocol was approved by the local Ethics Committee of Sao Paulo State University (#39844814.5.0000.5465).

### Consent to participate

All participants gave their written informed consent before they participated in the study.

### Consent for publication

All participants gave their written consent for publication.

### Declaration of conflicting interest

The author(s) declared no potential conflicts of interest with respect to the research, authorship, and/or publication of this article

### Funding statement

This research was supported by the Sao Paulo Research Foundation (FAPESP) [2014/22308- 0; 2017/19845-1], and the Coordenação de Aperfeiçoamento de Pessoal de Nível Superior— Brasil (CAPES)—Finance Code 001.

---

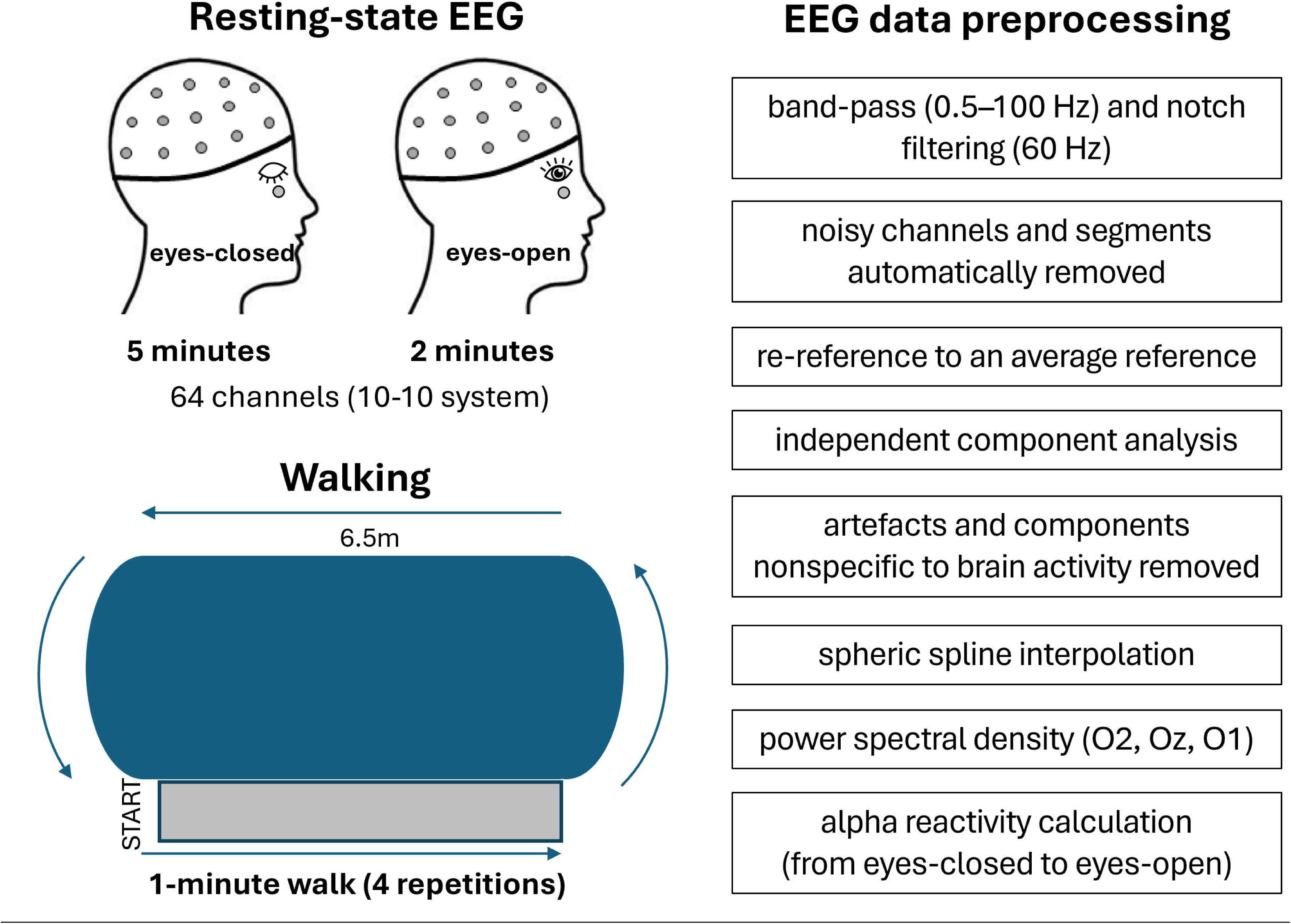

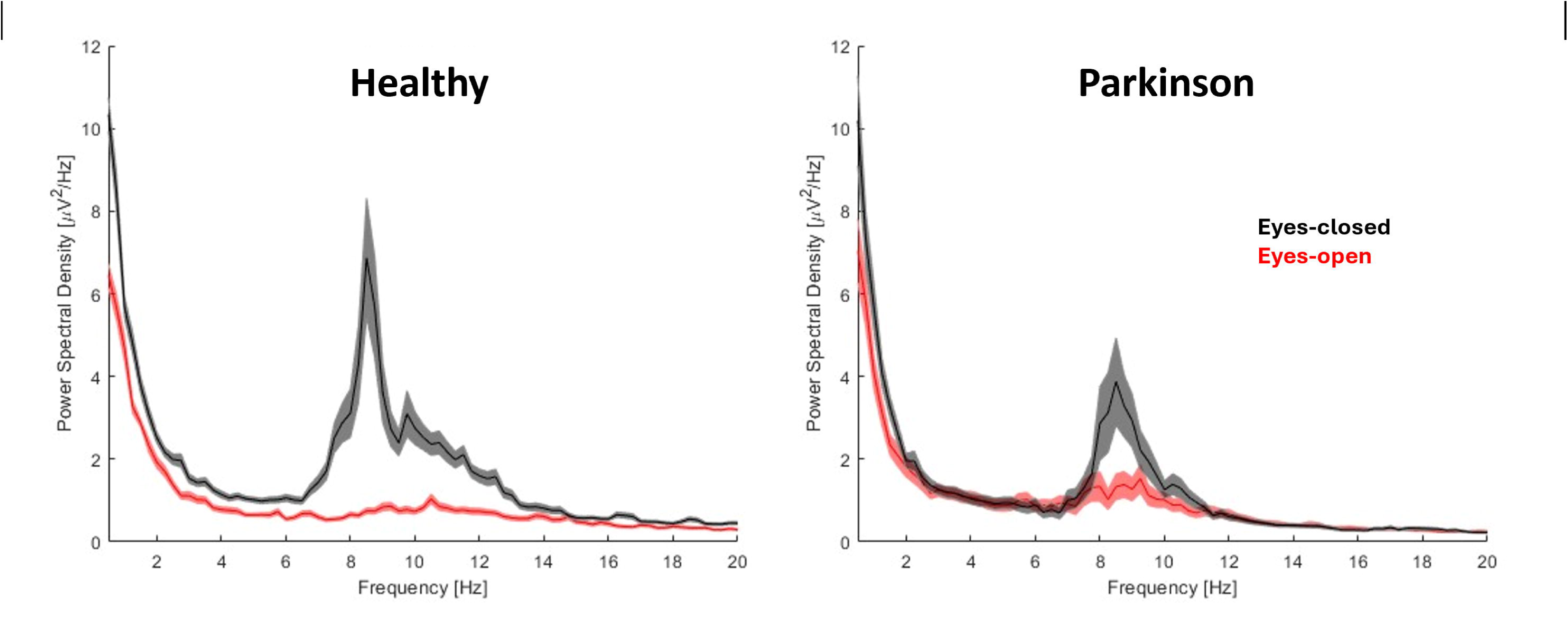

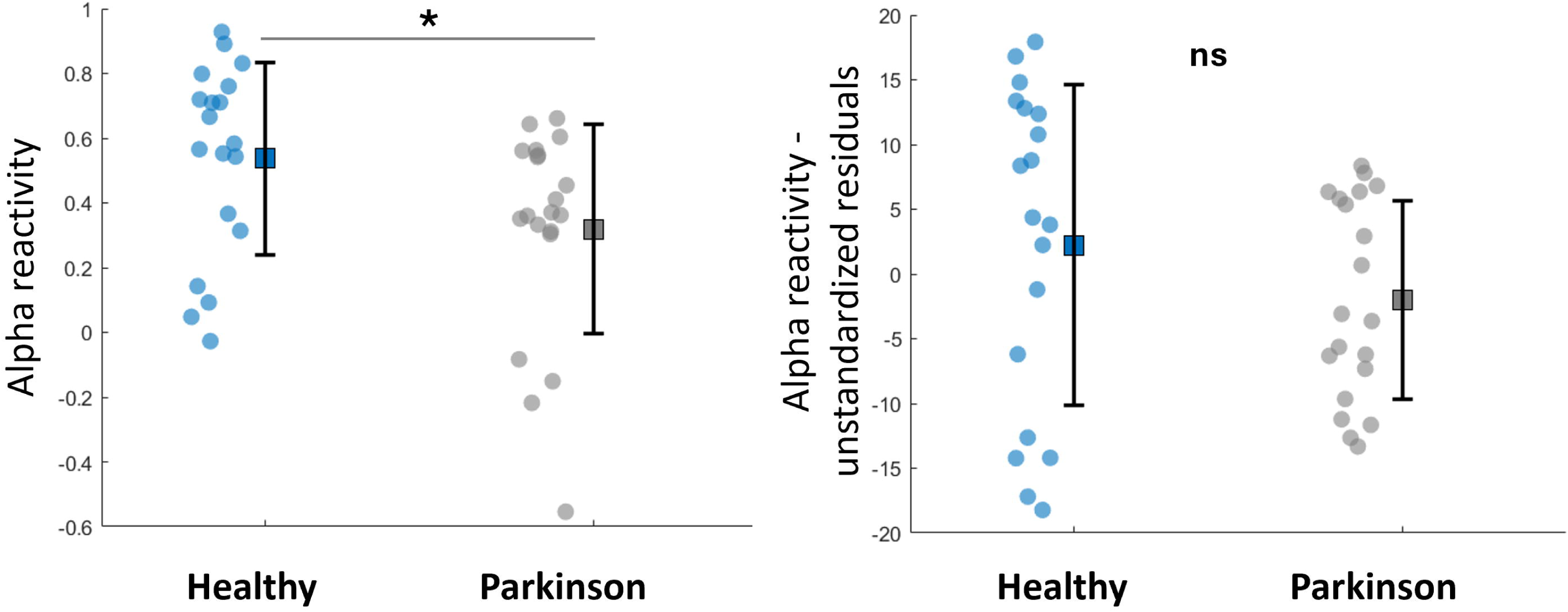

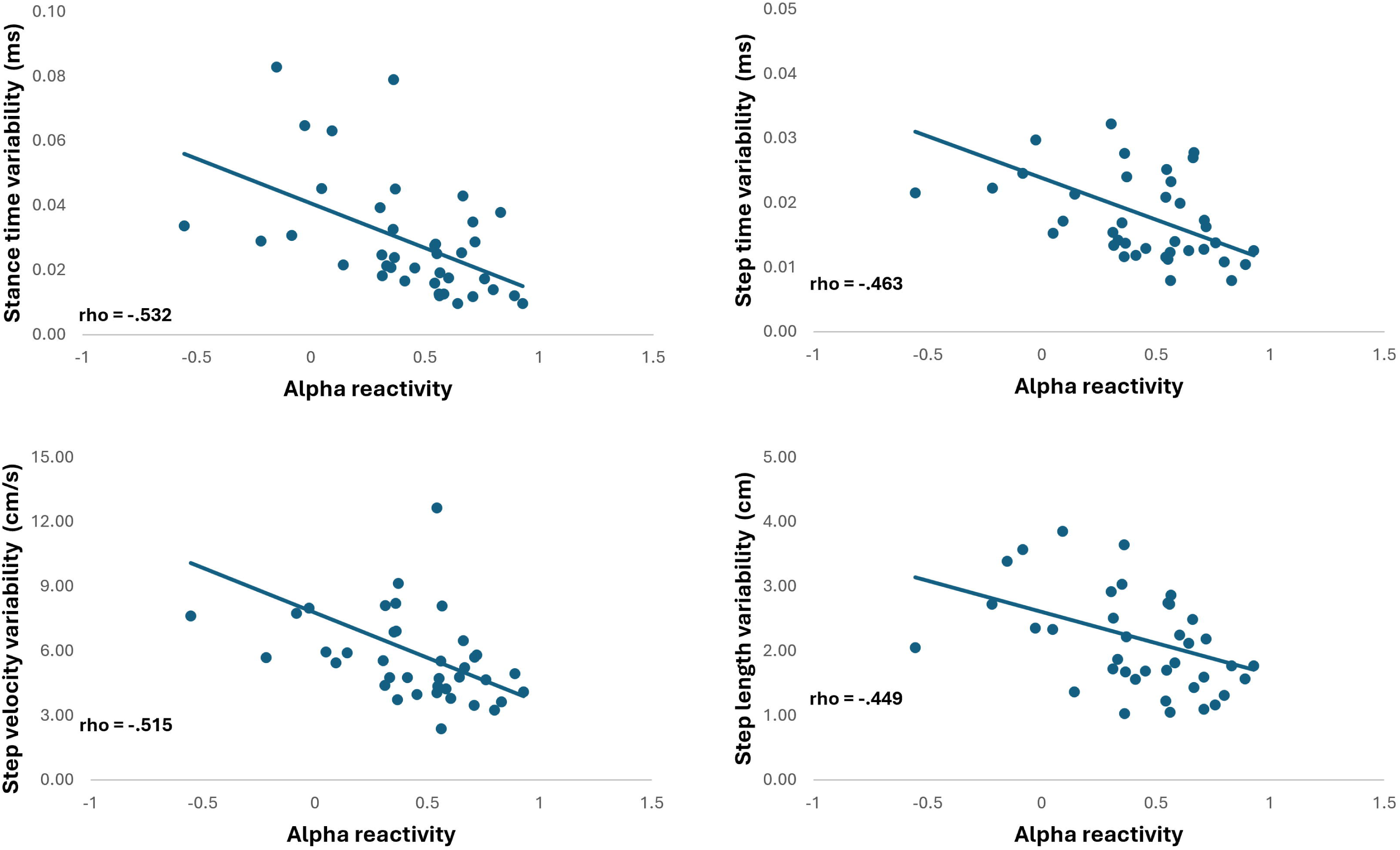

## Notes

**Conflict of interest**: The author(s) declared no potential conflicts of interest with respect to the research, authorship, and/or publication of this article.

### Competing Interest Statement

The authors have declared no competing interest.

