## Supplementary Table 1 for "Resting-state alpha reactivity is reduced in Parkinson’s disease and associated with gait variability"

**Supplementary Table 1.** Spatiotemporal gait measures for both groups.

| **Variables** | **Parkinson** | **Control** | **p-value** |
| --- | --- | --- | --- |
| Step velocity (cm/s) | 109.4 (19.0) | 120.5 (15.6) | 0.026* |
| Step length (cm) | 56.4 (9.0) | 63.7 (5.8) | 0.002* |
| Step time (ms) | 518.0 (26.6) | 532.4 (43.5) | 0.112 |
| Step width (cm) | 9.0 (3.5) | 8.5 (1.7) | 0.293 |
| Swing time (ms) | 379.6 (21.5) | 397.7 (27.7) | 0.014* |
| Stance time (ms) | 657.7 (42.2) | 665.7 (63.7) | 0.322 |
| Step velocity variability (cm/s) | 6.8 (3.8) | 5.2 (1.5) | 0.053 |
| Step length variability (cm) | 2.4 (1.0) | 1.9 (0.7) | 0.035* |
| Step time variability (ms) | 20.9 (10.4) | 15.8 (5.9) | 0.035* |
| Step width variability (cm) | 2.0 (0.6) | 2.0 (0.4) | 0.491 |
| Swing time variability (ms) | 23.2 (14.2) | 21.1 (13.6) | 0.316 |
| Stance time variability (ms) | 30.6 (19.4) | 27.4 (16.7)- | 0.291 |

*significant difference between groups.
